# Cerebral cortical structures linked to intelligence

**DOI:** 10.1101/2025.10.06.680808

**Authors:** Chun-Ju Chou, Mark Fiecas, Elisabetta C. del Re, Eero Vuoksimaa, Chi-Hua Chen

**Affiliations:** Department of Radiology, University of California, San Diego, CA, USA; Department of Bioengineering, University of California, San Diego, CA, USA; Division of Biostatistics, University of Minnesota School of Public Health, MN, USA; Department of Psychiatry, Harvard Medical School, Boston, MA, USA; Institute for Molecular Medicine Finland (FIMM), Helsinki Institute of Life Science, University of Helsinki, Helsinki, Finland

**Keywords:** Intelligence, Cortical morphology, Genetically-informed parcellation, Genome-wide association study, Mendelian randomization

## Abstract

Understanding the neural basis of intelligence in humans remains an ongoing scientific pursuit. Early studies with small samples identified potential regions but lacked consistency across findings. Recent large-scale magnetic resonance imaging (MRI) datasets, using intelligence measures focused on verbal-numerical reasoning, now offer more robust opportunities for discovery. In this study (N=11,289), we showed the dorsolateral prefrontal cortex as exhibiting the strongest effect size and a significant causal relationship with intelligence, where larger surface area predicts higher intelligence as revealed by Mendelian randomization analyses. Additional regions, including the orbitofrontal and temporal cortices, also showed causal links to intelligence. These regions are critical for working memory, executive function, and language. Reverse causality analyses further indicated that higher intelligence contributes to increased total surface area and greater cortical thickness in the perisylvian language region. Our findings replicate prior evidence of a bidirectional relationship between total surface area and intelligence and further offer novel insights into regional cortical associations and causal effect. Collectively, these findings support a polyregional cortical configuration of intelligence, highlighting the dorsolateral prefrontal cortex—a key hub for cognitive ability.

## Introduction

Genetic influences account for as high as 80% of the variance in individual differences in intelligence, also known as general cognitive ability (GCA) or the g-factor (1), yet its neural underpinnings remain unclear. In this study, we use the terms intelligence and GCA interchangeably. Over three decades of magnetic resonance imaging (MRI) studies have established a positive correlation of about 0.24 between brain volume and intelligence (2). In the past decade, studies (3) have shifted focus from brain volume to cortical surface area (SA) and cortical thickness (CT) – highly heritable, developmentally distinct and anatomically orthogonal traits (3). Evidence suggests the brain volume-intelligence link largely due to genetic effects and driven by total SA rather than mean CT (4).

Prior work indicates that GCA correlates with greater SA and thinner cortex in high-expanded regions (5, 6). More recent large-scale studies, such as those leveraging the UKB (7) and ABCD cohorts, provide valuable MRI data and have reported regional associations with intelligence (9–11). Other investigations have utilized GWAS summary statistics rather than individual-level data to examine brain–cognition relationships (12–14). Together, these studies showed global brain measures contribute to cognitive ability. However, the choice of cortical parcellation (e.g., number and boundaries of regions) and imaging metrics (e.g, volume or surface-based measures) impacts detection power and can influence the reproducibility and localization of associations (15). Consequently, although global brain measures show the strongest associations with intelligence, further research is needed to establish consistent evidence for regional contributions. Given the strong genetic influences on cortical structure and intelligence (4, 6), it is beneficial to consider the genetic nature of cortical regionalization (16, 17).

To address these gaps, we utilized a genetically defined cortical parcellation, which aligns with key functional specializations of the human cortex, to measure both SA and CT within each subdivision. Unlike traditional anatomical atlases, genetically-based parcellation delineates boundaries that are biologically grounded and genetically homogeneous (17). They further offer greater sensitivity for detecting regional associations (16). We aimed to identify the cortical correlates underlying intelligence in the UKB sample (N = 11,289). We hypothesized that genetically derived regions would reveal a polyregional neural substrate of intelligence, with SA and CT associations following distinct spatial patterns. Additionally, we expected to identify causal relationships between cortical structure and intelligence within this polyregional network through Mendelian Randomization analysis.

## Methods

### 1. Sample

The genomic, neuroimaging, cognitive, and demographic data were extracted from the UK Biobank (UKB) population cohort, under accession number 27412 (7, 18–20). Quality control (QC) procedures for imaging and demographic variables followed our previous work(16, 21). In this study, we included 34,842 participants with available neuroimaging. We accounted for genetic relatedness and removed related individuals from our analyses. Utilizing Genome-wide Complex Trait Analysis (GCTA) (22), the pairwise genetic relationship matrix (GRM) based on genome-wide autosomal variants was calculated, and 859 individuals were removed due to relatedness from pairs with an estimated GRM greater than 0.1 (indicating relatedness closer than third cousins), resulting in 33,861 participants. Excluding participants with missing cognitive scores, we resulted in a final dataset of 11289 participants (age range: 45.13-80.17 years, male-female ratio of 0.94) for subsequent analyses.

### 2. Genotype data

We used UKB Version 3 release of imputed genotype data and removed individuals with more than 10% missingness, as well as SNPs with more than 5% missingness, failing the Hardy-Weinberg equilibrium test at *p* = 1e-6 or with minor allele frequencies (MAF) below 0.01.

### 3. MRI data and atlases

T1-weighted MRI scans were collected from three scanning sites throughout the United Kingdom, all on identical Siemens Skyra 3T scanners (7). The MRI data processing steps were previously described in detail (16).

For cortical phenotypes, we adopted two genetically-informed atlases, including 12 regions for surface area and 12 for cortical thickness, and 2 global measures of total surface area and mean thickness. These atlases were developed using a data-driven fuzzy clustering technique to identify cortical parcels that are maximally genetically correlated, based on MRI scans from over 400 twins (17, 23). We combined measures of each phenotype across both hemispheres, in view of the largely bilateral symmetry of genetic patterning demonstrated previously (16, 24).

### 4. Fluid intelligence scores

GCA was operationalized as fluid intelligence scores that were extracted from the UK Biobank using field ID 20016. This is a simple unweighted sum of the number of correct answers given to the 13 fluid intelligence questions. Participants who did not answer all questions within the allotted 2-minute limit received a score of zero for each unattempted question. For this study, we included fluid intelligence scores from the first visit.

Intelligence was assessed using the UKB “fluid intelligence” score (Field ID 20016), which is more accurately described as a verbal-numerical reasoning (VNR) measure. Although this cognitive measure is labeled as assessing fluid cognitive abilities, which typically decline with aging, we note that its correlation with age in this UKB sample was very close to zero (-0.026).

This suggests that the score reflects both fluid and crystallized (stable) abilities. Indeed, the UKB fluid intelligence score has been reported to relate to crystallized abilities and is one of the UKB cognitive measures with the highest correlation to GCA (25).

### 5. Cortex-intelligence score association analysis

Prior to association analysis, we regressed out scanner site and a proxy of scan quality (FreeSurfer’s Euler number) (26) from each morphometric measure, resulting in *residualized absolute measurements*. In a parallel analysis of the regional measures, we also regressed out corresponding global measures (i.e. total surface area and mean cortical thickness for area and thickness phenotypes, respectively), referring to *residualized relative measurements*. This was done to study if the observed effects were specific to the region of interest, rather than due to the association with global measures. Subsequently, the residuals of each measure were scaled using rank-based inverse normal transformation to ensure normally distributed input. Because age can have nonlinear effects, we also compared models including a spline term for age (ns(Age, 3)) to linear age models. Results for associations between fluid intelligence scores and brain morphometric measures were essentially unchanged, so linear age was retained in the main analyses, with spline models noted as a sensitivity check.

Linear regression models examined associations between fluid intelligence scores and brain morphometric measures. Model 1: fluid intelligence scores as the dependent variable, with a cortical region’s residualized absolute measurement as the independent variable. Age, sex and diagnosis were covariates. Model 2: Same as Model 1, but using the cortical region’s residualized relative measurement. This approach allowed us to determine whether the observed effects were specific to the regions of interest rather than driven by global brain measures.

For the diagnosis covariate, we controlled for whether an individual had a brain-related diagnosis based on ICD10 diagnostic information collected by the UKB under field ID 41202. Individuals were classified as having a brain diagnosis if they met criteria for at least one class F (mental and behavioral disorders) or class G (disorders of the nervous system) diagnosis, with the exception of G56 - carpal tunnel syndrome, which is an extremely common condition and thus we did not consider it as a neurological diagnosis.

### 6. Determining effective number of independent phenotypes

To consider the potential correlation between phenotypes, we applied matSpD to determine the effective number of independent phenotypes (t_e_) (27), using correlation matrices of cortical measures. Statistical significance was then defined by Bonferroni correction for multiple comparisons (p < 0.05/t_e_).

In this study, we included regional measures (12 SA and 12 CT, averaged across hemispheres) and global measures (total SA and mean CT). To account for multiple comparisons, we calculated the effective number of independent traits as *t*_e_ = 22. Therefore, we set the study-wise significance threshold at p < 2.3e-3 (0.05/*t*_e_).

### 7. Mendelian randomization

We performed generalized summary-based MR (GSMR) (28) to evaluate the causal effects of the above 24 regional and 2 global morphometric measures on GCA, based on the summary statistics from previous genome-wide association analyses on the exposure (16) and outcome (29) phenotypes (detailed in Supplementary Methods 1, 2). These GWAS studies have non-overlapping samples. Linkage disequilibrium-independent (r^2^ < 0.05, 250kb) significant SFNPs (*p* < 5e-8) were included as candidate instrumental variables. The final set of instrumental variables used in the analyses is shown in Supplementary Information (Table S4, 5). Significance for GSMR analyses was defined as Bonferroni-corrected *p*-value < 2.3e-3.

## Results

### 1. Sample descriptive demographic

In this study sample, males were slightly older than females (64.54 ± 7.56 vs. 63.24 ± 7.32 years) but did not differ significantly in years of education (Table 1). Although males exhibited significantly higher intelligence scores, the effect size was small (Cohen’s d = -0.12). As expected, males had significantly greater SA than females (Cohen’s d = -1.27), while females had slightly thicker cortex (Cohen’s d = 0.27). The global brain differences align with males’ generally larger body size, as SA correlates with height. The thicker cortex in females may reflect the modest negative correlation between SA and CT observed here (r ≈ -0.1) and in prior studies (5, 6, 16, 30).

**Table 1.** Means and standard deviations by sex for demographic characteristics, intelligence measure and global brain metrics.

| Parameters | Group |  |  | Sex difference |  |
| --- | --- | --- | --- | --- | --- |
|  | Female<br>(n=5814) | Male<br>(n=5475) | Total | T-test<br>p-value | Cohen's d |
| Age (yrs) | 63.24 ± 7.32 | 64.54 ± 7.56 | 63.87 ± 7.46 | < 1e-10 | -0.18 |
| Education<br>(final age) | 17.10 ± 2.39 | 17.17 ± 2.84 | 17.13 ± 2.61 | 0.31 | -0.03 |
| GCA | 6.71 ± 1.93 | 6.94 ± 2.06 | 6.82 ± 2.00 | 3.41e-10 | -0.12 |
| Total SA (cm<sup>2</sup>) | 1605.86 ± 126.70 | 1776.58 ± 141.60 | 1688.66 ± 158.97 | < 1e-10 | -1.27 |
| Mean CT (mm) | 2.43 ± 0.09 | 2.40 ± 0.10 | 2.41 ± 0.10 | < 1e-10 | 0.27 |

Note: Data is presented as mean ± standard deviation (SD). GCA = general cognitive ability as measured with UKB fluid intelligence score, SA = cortical surface area, CT = cortical thickness. P-values from the t-test for sex differences. In Cohen's d, negative values indicate greater values in males

### 2. Absolute and relative cortical morphology associated with intelligence

Two regression models were performed to examine the cortical-intelligence associations: one unadjusted and one adjusted for overall brain size (total SA or mean CT), corresponding to absolute and relative measures, respectively.

When analyzing absolute cortical morphology, we confirmed a strong positive association between total SA and intelligence (β=0.34, p=<1e-10, R^2^=2.7%). For regional SA, all cortical regions displayed significant positive associations with intelligence (Figure 1, Table S1). The dorsolateral prefrontal area, subserving working memory and executive functions, exhibited the highest effect size (β = 0.35; R^2^ = 2.8%) among all regions, slightly surpassing total SA. Other top regions included the superior temporal and orbitofrontal areas, which are critical for auditory/language processing and emotional working memory, respectively. In contrast to the SA results, mean CT showed a nominally significant positive association (β=0.06, p=7.36e-03) and regional CT exhibited generally modest associations with intelligence (Figure 1, Table S1).

**Figure 1.**
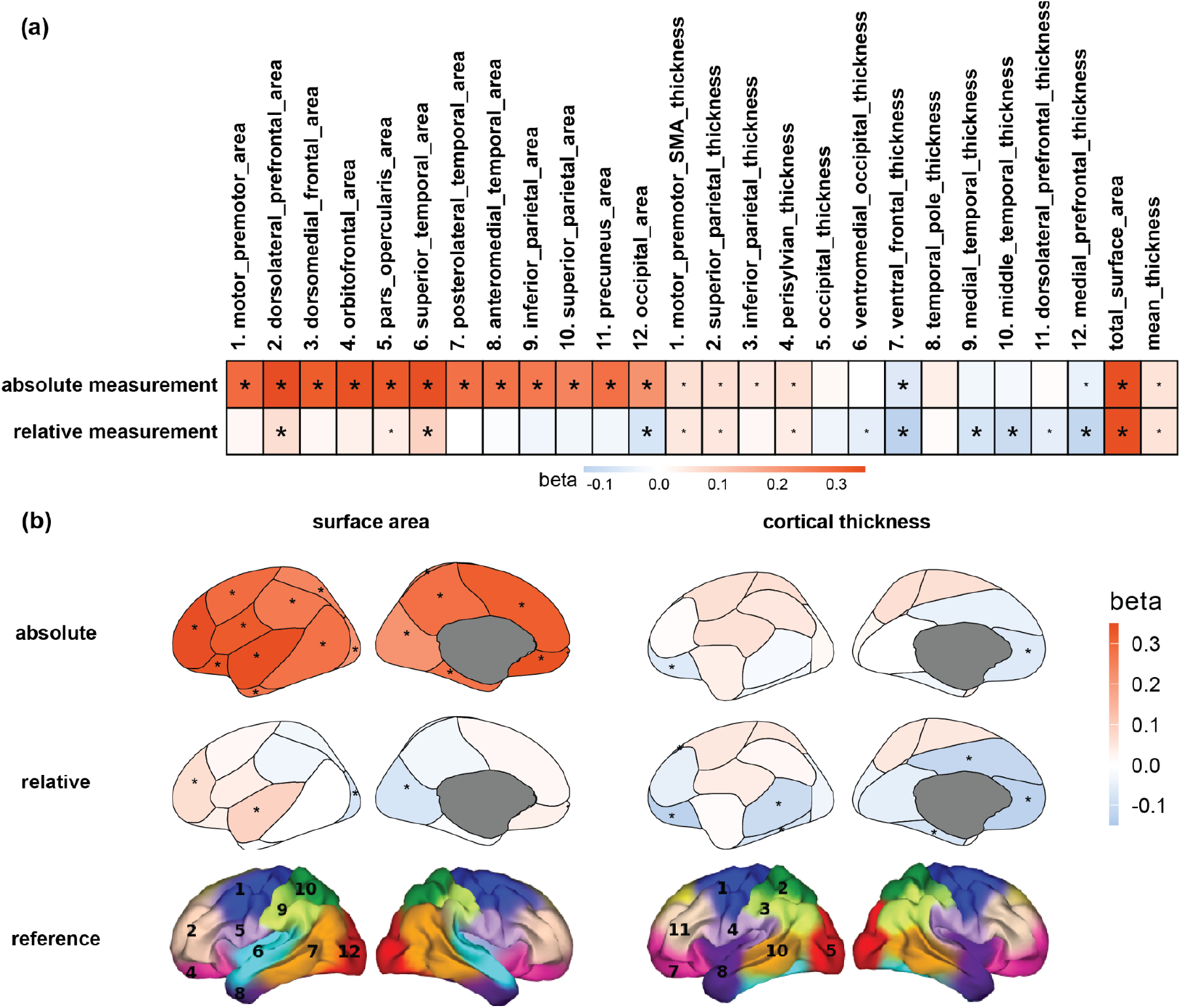
(a) Association between cortical morphology and intelligence, with and without adjustment for global measures. Big or small asterisks denote statistical (*p* < 2.3e-3) or nominal (*p* < 0.05) significance. Global measures are presented twice at the end of both rows of heatmaps. (b) Brain maps with color shading represent beta coefficients for surface area and cortical thickness. Asterisks indicate cortical regions with significant associations after correction for multiple comparisons (*p* < 2.3e-3). The bottom row shows atlas brain maps for reference, with numbering corresponding to the regions listed above the heatmap.

After adjusting for total SA, relative cortical SA showed smaller associations. However, the superior temporal area (β=0.09, p=2.05e-06) and dorsolateral prefrontal area (β=0.07, p=4.98e-04) remained positively associated with intelligence, whereas the occipital area showed a significant negative association (β=-0.08, p=5.24e-05). For CT, negative associations for relative CT were observed after adjusting for mean CT, particularly in ventral frontal (β=-0.13, p=1.97e-11), middle temporal (β=-0.10, p=1.11e-07), and medial prefrontal (β=-0.11, p=1.53e-09) regions.

### 3. Bidirectional causal relationship for cortical morphology and intelligence

Beyond associations, we next studied the bidirectional causal relationships between cortical structure and intelligence to better contextualize these links (Figure 2, Table S3). In forward MR analyses, total SA exhibited a causal effect on intelligence (βxy=0.13, CI = [0.10,0.18], p=2.08e-10). At the regional level, larger dorsolateral prefrontal (βxy=0.09, CI = [0.02,0.15], p=1.05e-02), posterolateral temporal (βxy=0.06, CI = [0.02,0.10], p=4.00e-03), and orbitofrontal (βxy=0.06, CI = [0.00,0.13], p=4.19e-02) areas had positive causal effects on intelligence, whereas the superior parietal area had a negative causal effect (Figure 2). For CT, no global or regional CT measures demonstrated a causal influence on intelligence.

**Figure 2.**
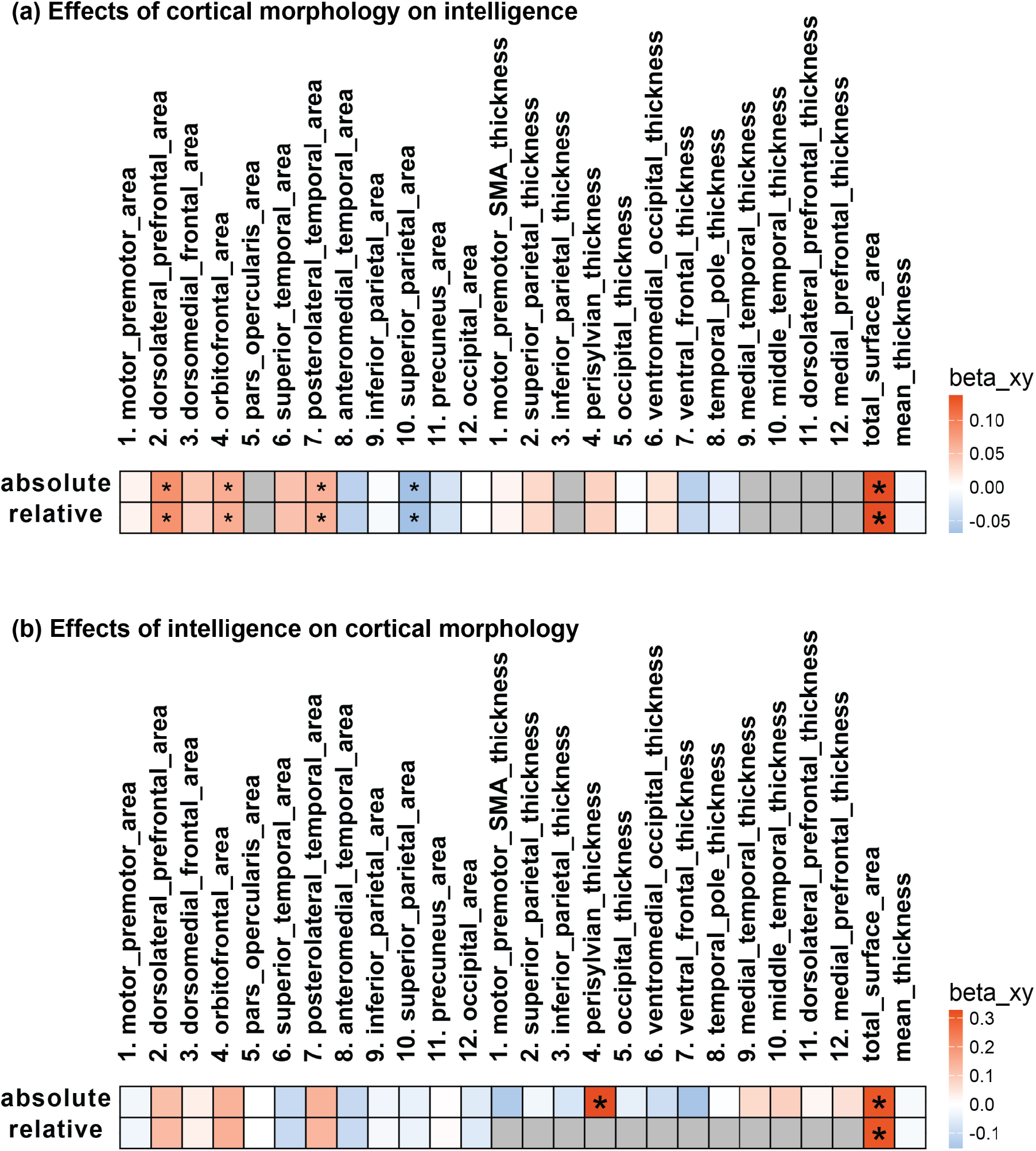
Bidirectional causal effects between cortical morphology and intelligence. The heat maps display both (a) forward and (b) reverse Mendelian Randomization analyses. Big or small asterisks denote statistical (*p* < 2.3e-3, Bonferroni correction) or nominal (*p* < 0.05) significance. Gray boxes indicate missing data due to limitations such as restricted sample size or insufficient number of filtered single nucleotide polymorphisms (SNPs) meeting the inclusion threshold.

In reverse MR analyses, intelligence was found to causally influence total SA (βxy=0.31, CI = [0.12,0.49], p=1.02e-03), suggesting a reciprocal relationship. At the regional level, higher intelligence causally influenced perisylvian thickness (βxy=0.33, CI = [0.14,0.51], p=5.19e-04).

## Discussion

Our study replicated the finding that total SA, rather than mean CT, is linked to intelligence (4, 30), and further provided novel insights into regional cortical associations with intelligence. Adjusting for global SA reduced the strength of regional associations, consistent with previous findings (6, 31). Nevertheless, we found that the superior temporal and dorsolateral prefrontal areas remained positively associated with intelligence, even after accounting for global SA. Prior work has shown that global SA has the highest genetic correlation with dorsolateral prefrontal SA, suggesting that adjusting for global measures may substantially attenuate regional associations (21). The persistence of these associations may reflect their robustness.

We observed an anterior-posterior gradient in SA-intelligence associations, with the strongest effects in the dorsolateral prefrontal and superior temporal regions, including the insula and Sylvian fissure (17, 32). In contrast, CT-intelligence associations, though more modest, indicated a pattern of a dorsal-ventral gradient. These patterns are consistent with those reported in both our previous research and other studies of cortical patterning (32). As shown here for the first time in relation to intelligence, we propose that these gradients may reflect crucial aspects of cortical architecture and cognitive functioning. A prior study using the same genetic parcellation also found the strongest associations in the prefrontal and temporal cortices, though only based on unadjusted SA. This suggests that these findings are robust and replicated across independent samples and age groups (5).

Mendelian Randomization (MR) analyses provided evidence of causality between cortical structure and intelligence. The bidirectional relationship between total SA and intelligence was also reported in a recent study (33), suggesting a dynamic interplay between cortical structure and cognitive function. Here we provided further insight into regional relationships. Notably, regional SA in prefrontal and temporal areas exerted a unidirectional causal effect on intelligence. Note that the dorsolateral prefrontal and posterolateral temporal areas were part of the “high-expanded” regions identified previously (5, 6), highlighting their genetic correlations with GCA (6). Whereas intelligence influenced perisylvian thickness, a region with a known role in language processing, possibly reflecting experience-dependent cortical plasticity (34). One caveat is that the statistical power of the MR analysis was limited for some regions, preventing us from yielding results across all phenotypes.

Lin et al. (2025) used the Desikan atlas, which does not delineate the lateral prefrontal cortex directly but only proxy regions such as the rostral and caudal middle frontal gyri.(14) In contrast, our genetically informed parcellation delineates the dorsolateral prefrontal cortex itself, which we identified as the most significant region in MR analyses of intelligence—consistent with long-standing evidence for its central role in cognitive ability. Lin et al. instead reported the insula and transverse temporal cortex as their top regions associated with cognition, areas that may contribute but are not primary loci emphasized in the Parieto-Frontal Integration Theory (P-FIT) (35–38). Methodologically, Lin et al. used a more liberal p-value threshold for instrument selection to boost power (p < 10^-6^ p or 10^-7^), while we applied the stricter genome-wide significance threshold (p < 5 × 10^-8^) for more conservative inference.

Our reverse MR analyses suggested that intelligence may influence cortical surface area, consistent with prior reports of bidirectional causal links (12, 14, 33). Similarly, Schnack et al. showed that longitudinal changes in cortical thickness and surface area are correlated with intelligence over time, supporting the role of developmental cortical remodeling and experience-dependent plasticity throughout life (39).

The present study underscores the significance of the dorsolateral prefrontal cortex, a key region for higher-order cognition such as working memory and executive functions, as demonstrated by evidence from brain lesion patients (11, 40). We identified this region as having the strongest effect size and a significant causal relationship with intelligence. Other key regions identified here include the orbitofrontal and temporal cortices. In contrast, we found limited support for significant parietal cortex associations, aligning only partially with P-FIT (35–38). Similarly, a UKB study (11) using absolute volumetric measures and g-factor based on four cognitive tests reported weaker parietal involvement (11, 40), though they also identified occipital contributions, which we did not observe. These discrepancies may be attributed to differences in study design in that our study used SA and CT instead of volume, investigated regional effects adjusted for global cortical size, and included verbal-numerical reasoning as the measure of intelligence.

In conclusion, these findings add to the ongoing discussions about the neural basis of intelligence. The results align with previous research (6), showing that higher intelligence is linked to expanded cortical SA in key association cortical areas important for higher cognitive function. Our results support a polyregional model that includes the prefrontal and temporal cortices important for working memory, executive functions and language. Moreover, we demonstrated causal relationships between these regions and intelligence, further underscoring their significance.

**Acknowledgement**

This research was supported by the National Institute of Mental Health under R01MH132783. E. Vuoksimaa was supported by the Sigrid Jusélius Foundation and the Research Council of Finland (grants 314639 and 345988). This research has been conducted using data from the UK Biobank, a major biomedical database, under application number 27412.

## Data Availability

This study utilizes individual-level genetic and imaging data from the UK Biobank (https://www.ukbiobank.ac.uk/). The genome-wide association data for cortical regions were provided from our previously published studies, which can be accessed via the GWAS Catalog (https://www.ebi.ac.uk/gwas/publications/35113692 and https://www.ebi.ac.uk/gwas/publications/36893272). The genome-wide association data for intelligence were obtained directly from the authors upon request, as a means of avoiding overlap with UKB samples, and the original source was from the PGC (https://pgc.unc.edu/) or the GWAS Catalog (https://www.ebi.ac.uk/gwas/publications/29942086). GSMR method and code are publicly available via the GCTA software repository on GitHub (https://github.com/JianYang-Lab/gsmr/releases).

