## Supplementary Information for "Cerebral cortical structures linked to intelligence"

### Table of Contents

**Supplementary Methods.**
**Supplementary Results.**
**SFigure 1.**
**Supplementary Table.**
**Supplementary Reference.**

**Supplementary Methods.**

1. Details of Mendelian Randomization Analyses

To evaluate the robustness of our causal inference and address potential violations of MR assumptions, we implemented five complementary MR approaches: GSMR (1), inverse-variance weighted (IVW), MR-Egger regression, weighted median, and weighted mode (2–4). Each method relies on different assumptions and offers different sensitivity to pleiotropy or invalid instruments.

For GSMR, SNP instruments were first selected from the exposure GWAS at genome-wide significance ($p$ < 5e-8) and pruned for linkage disequilibrium ($r^2 < 0.05$, 250kb). For each retained SNP, we calculated instrument strength using the F-statistic, with all instruments exceeding the conventional threshold of $F > 10$, ensuring sufficient strength. While Mendelian Randomization guidelines suggest a rule of thumb of at least 10 independent instruments for robust sensitivity analyses, prior work has emphasized that analyses with fewer variants can still yield valid inference if the instruments are strong and results are interpreted cautiously (5, 6). This applied to our reverse MR models, where fewer than 10 instruments were available but all exhibited strong F-statistics. GSMR applies the HEIDI-outlier test (pHEIDI < 0.01) to identify and remove individual SNPs whose outcome effects deviate from expectation, thereby filtering potential pleiotropic variants before causal estimation. While this approach directly addresses exclusion restriction violations, residual pleiotropy may persist if effects are small or widespread. IVW provides high statistical power by combining all instruments under the assumption that they are valid or that pleiotropic effects are balanced on average; however, it is vulnerable to directional pleiotropy. MR-Egger regression relaxes this assumption by allowing for a non-zero intercept, which can detect and adjust for directional pleiotropy, but requires the InSIDE assumption and has lower power with wider confidence intervals. The weighted median estimator yields consistent causal effects if at least 50% of the total instrument weight comes from valid SNPs, offering robustness to invalid instruments at the cost of some efficiency. Finally, the weighted mode method identifies the most common causal effect across instruments and assumes that the largest cluster represents valid instruments, providing robustness in cases where only a subset of SNPs are valid but with reduced statistical power. By aligning the results across these complementary methods, we were able to assess the stability of causal estimates under different model assumptions and better evaluate the potential impact of assumption violations.

2. Details of GSMR exposure and outcome

For the exposures, we used genome-wide association results of cortical morphology derived from MRI-based measures of cortical surface area and thickness in the UK Biobank imaging cohort (N ≈ 33,000 after quality control) (7, 8). For the outcome, we used publicly available summary statistics from the large-scale genome-wide association study of intelligence reported by Savage et al (9). This meta-analysis included approximately 270,000 individuals of European ancestry from multiple cohorts (e.g., UKB, COGENT, RS, STR, TEDS). Intelligence was measured through harmonized cognitive performance assessments across contributing studies. Different cognitive measures were harmonized to index a common latent *g* factor. To avoid sample overlap with our UKB brain morphology GWAS, we specifically requested and used the UKB-excluded summary statistics provided by the authors, which were based on N = 74,214 individuals across 13 non-UKB cohorts. These final exposure (33,181) and outcome (74,214) sample sizes represent the non-overlapping datasets used for our GSMR analyses.

By combining these GWAS datasets, we implemented GSMR to test the causal relationship between genetically predicted cortical morphology and intelligence, while minimizing bias due to population stratification, relatedness, or sample overlap.

3. Details of matSpD Determining Methods

The *matSpD* software uses matrix spectral decomposition to account for correlations among phenotypes or genetic markers in multiple testing corrections. By applying eigen decomposition to the correlation matrix, it estimates the effective number of independent variables (*te*) (10).

Instead of treating correlated phenotypes as fully independent, *matSpD* uses the variance explained by eigenvalues to determine how many variables contribute unique information. This effective number is then used to adjust the Bonferroni threshold, providing a less conservative correction while controlling the family-wise error rate.

In this study, we applied *matSpD* to cortical and subcortical correlation matrices to estimate *te*, which defined the Bonferroni-corrected significance threshold ($p < 0.05/te$).

**Supplementary Results.**

1. Bidirectional causal effects between cortical morphology and intelligence with different MR methods comparison

To avoid potential MR assumption violations, we complemented GSMR with additional MR methods. From Supplementary Figure 1, we observe overlapping significance across approaches in total surface area and the posterolateral temporal area, with forward MR results aligning across both absolute and relative analyses. (Table S2) For the dorsolateral prefrontal region, GSMR detected significance that was also supported by MR-IVW. Other sensitivity methods did not reach significance for this region, which may reflect reduced power under these more conservative models.

### Supplementary Figures.

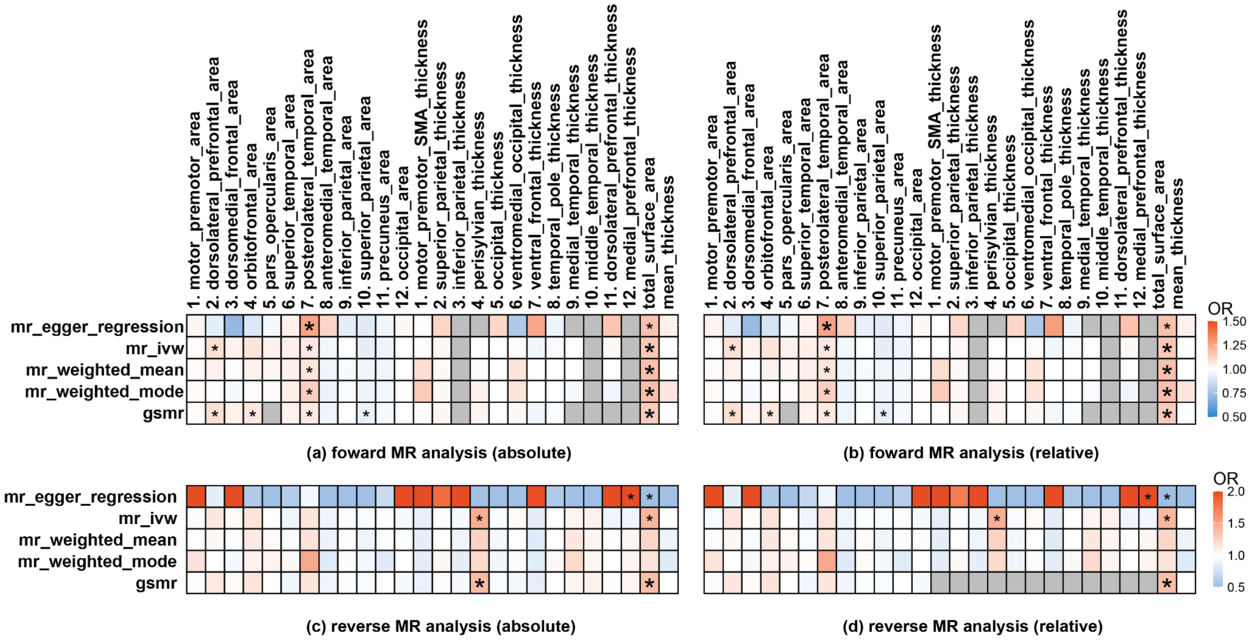

**Figure S1.** Bidirectional causal effects between cortical morphology and intelligence with different MR methods comparison. The heat maps display both (a)(b) forward and (c)(d) reverse Mendelian Randomization analyses. Big or small asterisks denote statistical ($p$ < 2.3e-3, Bonferroni correction) or nominal ($p$ < 0.05) significance. Gray boxes indicate missing data due to limitations such as restricted sample size or insufficient number of filtered single nucleotide polymorphisms (SNPs) meeting the inclusion threshold.

**Supplementary Table.**

Table S4. Instrument strength of genome-wide significant SNPs used in forward GSMR analyses across absolute cortical regions after pleiotropy filtering.

| Forward GSMR IV (Significant Regions) | | | | | | | | |
|---|---|---|---|---|---|---|---|---|
| Dorsolateral prefrontal area | | | Orbitofrontal area | | | Posterolateral temporal area | | |
| IVs | F-stat | p | IVs | F-stat | p | IVs | F-stat | p |
| rs1015446 | 34.9401 | 3.40E-09 | rs1015446 | 67.0930 | 2.59E-16 | rs1023470 | 31.2423 | 2.28E-08 |
| rs1078363 | 31.1565 | 2.38E-08 | rs10206105 | 32.1853 | 1.40E-08 | rs10809913 | 37.7563 | 8.02E-10 |
| rs11020063 | 43.7425 | 3.75E-11 | rs10942525 | 33.6054 | 6.75E-09 | rs10830941 | 36.9991 | 1.18E-09 |
| rs12676765 | 31.4399 | 2.06E-08 | rs11579220 | 38.4984 | 5.48E-10 | rs10878269 | 93.4330 | 4.20E-22 |
| rs1456979 | 39.2943 | 3.64E-10 | rs11767221 | 78.7286 | 7.13E-19 | rs114921579 | 35.1844 | 3.00E-09 |
| rs17235841 | 38.7386 | 4.85E-10 | rs12626264 | 30.6217 | 3.14E-08 | rs11669334 | 33.3128 | 7.85E-09 |
| rs4868084 | 51.0033 | 9.22E-13 | rs12724488 | 42.7068 | 6.36E-11 | rs12372133 | 41.2045 | 1.37E-10 |
| rs4924346 | 49.0254 | 2.53E-12 | rs13233151 | 43.0543 | 5.32E-11 | rs1637571 | 36.7538 | 1.34E-09 |
| rs55743972 | 38.8828 | 4.50E-10 | rs2282042 | 32.2479 | 1.36E-08 | rs2166123 | 36.5849 | 1.46E-09 |
| rs7093914 | 32.0145 | 1.53E-08 | rs4681356 | 37.0855 | 1.13E-09 | rs2991716 | 30.8946 | 2.72E-08 |
| rs888812 | 63.4937 | 1.61E-15 | rs7451690 | 33.6932 | 6.45E-09 | rs305448 | 31.3495 | 2.16E-08 |
| | | | rs9601373 | 31.3759 | 2.13E-08 | rs3940842 | 55.6570 | 8.63E-14 |
| | | | | | | rs3999543 | 33.6925 | 6.46E-09 |
| | | | | | | rs61741508 | 49.1097 | 2.42E-12 |
| | | | | | | rs61909999 | 32.6133 | 1.12E-08 |
| | | | | | | rs669952 | 109.6043 | 1.20E-25 |
| | | | | | | rs68175985 | 130.1150 | 3.87E-30 |
| | | | | | | rs7154212 | 35.4162 | 2.66E-09 |
| | | | | | | rs7180955 | 30.5588 | 3.24E-08 |
| | | | | | | rs743029 | 30.9718 | 2.62E-08 |
| | | | | | | rs7560411 | 64.3856 | 1.02E-15 |
| | | | | | | rs7584720 | 43.4281 | 4.40E-11 |
| | | | | | | rs76014677 | 42.1546 | 8.43E-11 |
| | | | | | | rs79272390 | 38.5944 | 5.22E-10 |
| | | | | | | rs8025699 | 30.4096 | 3.50E-08 |
| | | | | | | rs8077219 | 38.0424 | 6.92E-10 |

| Superior parietal area | | | Total surface area | | |
|---|---|---|---|---|---|
| IVs | F-stat | p | IVs | F-stat | p |
| rs10496078 | 44.1771 | 3.00E-11 | rs10460587 | 46.6409 | 8.53E-12 |
| rs11175755 | 51.1470 | 8.57E-13 | rs10496091 | 34.7754 | 3.70E-09 |
| rs11669334 | 63.1193 | 1.95E-15 | rs11079849 | 36.9575 | 1.21E-09 |
| rs11979208 | 37.3260 | 9.99E-10 | rs111379945 | 46.3455 | 9.91E-12 |
| rs12615194 | 38.4811 | 5.53E-10 | rs11938781 | 41.1253 | 1.43E-10 |
| rs2033939 | 75.7102 | 3.28E-18 | rs139590428 | 31.7002 | 1.80E-08 |
| rs231267 | 33.4160 | 7.44E-09 | rs1951120 | 35.3004 | 2.83E-09 |
| rs4437022 | 31.4212 | 2.08E-08 | rs1979477 | 30.0570 | 4.20E-08 |
| rs56111638 | 59.7040 | 1.10E-14 | rs199534 | 226.5905 | 3.31E-51 |
| rs57205040 | 48.9207 | 2.67E-12 | rs2764264 | 52.7473 | 3.79E-13 |
| rs62109937 | 29.9630 | 4.40E-08 | rs28741121 | 31.0707 | 2.49E-08 |
| rs6858566 | 70.7845 | 3.98E-17 | rs3786824 | 35.3381 | 2.77E-09 |
| rs73006822 | 125.2955 | 4.39E-29 | rs4273712 | 121.2043 | 3.45E-28 |
| rs73578186 | 34.9155 | 3.44E-09 | rs4685 | 51.4943 | 7.18E-13 |
| rs7563957 | 57.8581 | 2.82E-14 | rs62283132 | 69.1649 | 9.06E-17 |
| rs888812 | 41.8035 | 1.01E-10 | rs7220421 | 70.2833 | 5.14E-17 |
| rs9345124 | 33.0605 | 8.93E-09 | rs76928645 | 83.5473 | 6.22E-20 |
| | | | rs7715167 | 33.0316 | 9.07E-09 |
| | | | rs77324770 | 36.3944 | 1.61E-09 |
| | | | rs8756 | 60.1247 | 8.90E-15 |
| | | | rs9266618 | 47.4516 | 5.64E-12 |

*Note: All listed SNPs exceed the conventional F > 10 threshold, indicating strong instruments. p-values reflect SNP–exposure associations from the discovery cortical GWAS.*

Table S5. Instrument strength of genome-wide significant SNPs used in reverse GSMR analyses across absolute cortical regions after pleiotropy filtering.

| Reverse GSMR IV (Significant Regions) | | | | | |
|---|---|---|---|---|---|
| Perisylvian thickness | | | Total surface area | | |
| IVs | F-stat | p | IVs | F-stat | p |
| rs12602130 | 29.7462 | 4.94E-08 | rs12602130 | 29.7462 | 4.94E-08 |
| rs2251499 | 32.9820 | 9.31E-09 | rs2251499 | 32.9820 | 9.31E-09 |
| rs2726491 | 34.3161 | 4.69E-09 | rs2726491 | 34.3161 | 4.69E-09 |
| rs2920940 | 29.9647 | 4.39E-08 | rs2920940 | 29.9647 | 4.39E-08 |
| rs4839715 | 33.1315 | 8.59E-09 | rs4839715 | 33.1315 | 8.59E-09 |
| rs6023759 | 30.9693 | 2.62E-08 | rs6023759 | 30.9693 | 2.62E-08 |
| rs7359397 | 30.9692 | 2.62E-08 | rs7359397 | 30.9692 | 2.62E-08 |
| rs7468585 | 35.8442 | 2.14E-09 | rs7468585 | 35.8442 | 2.14E-09 |

*Note: All listed SNPs exceed the conventional $F > 10$ threshold, indicating strong instruments. p-values reflect SNP–exposure associations from the discovery IQ GWAS.*

**Supplementary Reference.**
